# A comprehensive map of genetic variation in the world’s largest ethnic group - Han Chinese

**DOI:** 10.1101/162982

**Authors:** Charleston W. K. Chiang, Serghei Mangul, Christopher R. Robles, Warren W. Kretzschmar, Na Cai, Kenneth S. Kendler, Sriram Sankararam, Jonathan Flint

## Abstract

As are most non-European populations around the globe, the Han Chinese are relatively understudied in population and medical genetics studies. From low-coverage whole-genome sequencing of 11,670 Han Chinese women we present a catalog of 25,057,223 variants, including 548,401 novel variants that are seen at least 10 times in our dataset. Individuals from our study come from 19 out of 22 provinces across China, allowing us to study population structure, genetic ancestry, and local adaptation in Han Chinese. We identify previously unrecognized population structure along the East-West axis of China and report unique signals of admixture across geographical space, such as European influences among the Northwestern provinces of China. Finally, we identified a number of highly differentiated loci, indicative of local adaptation in the Han Chinese. In particular, we detected extreme differentiation among the Han Chinese at MTHFR, ADH7, and FADS loci, suggesting that these loci may not be specifically selected in Tibetan and Inuit populations as previously suggested. On the other hand, we find that Neandertal ancestry does not vary significantly across the provinces, consistent with admixture prior to the dispersal of modern Han Chinese. Furthermore, contrary to a previous report, Neandertal ancestry does not explain a significant amount of heritability in depression. Our findings provide the largest genetic data set so far made available for Han Chinese and provide insights into the history and population structure of the world’s largest ethnic group.

## Introduction

Over the last decade genome wide association studies (GWAS) and, more recently, sequencing studies have focused on examining the distribution of genetic variants in European populations, providing datasets to explore fundamental genetic questions in these populations. However comprehensive analyses require more complete characterization of non-European populations than is currently available. Here we describe a resource of genetic variants for the world’s largest ethnic group, Han Chinese.

To date, a range of strategies has been employed to characterize populations. In Genomes of the Netherlands (GoNL) (Genome of the Netherlands 2014), high quality genotypes at both single nucleotide and structural variations can be detected with intermediate (∼13X) coverage with higher than 99% accuracy, and the trio design enabled the investigation of de novo mutations. In a Sardinian (Sidore et al. 2015) and a Icelandic (Gudbjartsson et al. 2015) population cohort, extensive haplotype sharing within populations were used to inform accurate genotype calling among low (∼4-6X) and intermediate (∼20X) coverage sequencing of ∼2000-3000 individuals. In the UK10K project (UK10K et al. 2015), low (∼7x) whole-genome sequencing in 3,781 healthy samples from two British cohorts is combined with deep (80x) exome sequencing in three disease cohorts (rare diseases, severe obesity and neurodevelopmental disorders) and was able to accurately detect low frequency and rare variants associated with quantitative traits.

We adopt a different approach for the characterization of the Han Chinese. We obtained genetic variants from low coverage whole genome sequence information of 11,670 Han Chinese during a case control study of major depressive disorder (MDD) from the China, Oxford and Virginia Commonwealth University Experimental Research on Genetic Epidemiology (CONVERGE). With a median coverage of 1.7X our sequence data are predicted to identify rare (<0.5%) single nucleotide polymorphisms (SNPs) with high confidence (Li et al. 2011b) and obtain accurate estimates of allele frequencies in a large sample. Using this genomic resource, we catalog variants, and use the data to characterize the genetic structure of the Chinese population.

Our understanding of the population structure and history of the Han Chinese has been relatively limited. Historical records suggested that the Han Chinese originated from the Central Plain region of China during the early historic and pre-historic era. From there, because of their advantage in agriculture and technology the group expanded both northward and southward to become the largest ethnic group today in China (Zhao et al. 2015). This historical movement is corroborated genetically based on uniparental markers from both modern and ancient DNA samples (Wen et al. 2004; Zhao et al. 2015). Clear North to South structure is also evident from array-based genome-wide data (Chen et al. 2009; Xu et al. 2009; Chen et al. 2016), strongly driving the pattern in the first component of a principal components analysis (PCA). However, very little structure has been observed beyond the first principal component. This could be due to a lack of representative samples across China, and/or the small sample sizes that reduce the power to detect less prominent features of population structure. Furthermore, because typical representatives of Han Chinese are from the 1000 Genomes or HGDP data, where the Han Chinese are only divided into Northern and Southern Han Chinese in small number, little has been discussed of the relationship between Han Chinese and other, non-East Asian populations. Nonetheless, in these relatively limited reference samples of Han Chinese, elevated ancestry from Western Eurasians had been detected among Northern Han Chinese, and speculated to be due to trades along the Silk Road (Hellenthal et al. 2014).

Taking advantage of the diverse geographical sampling of Han Chinese across China in CONVERGE, covering 19 out of 22 provinces in China with hundreds of samples typically available per province, we sought to investigate the genetic structure of modern Han Chinese. Furthermore, using largely allele-frequency based population genetic analyses, we examine pattern of ancestry and admixture across China, and investigate regions of genome undergoing extreme differentiation in Han Chinese. We also investigate the relationship of modern Han Chinese with ancient and archaic samples. As East Asians are thought to possess archaic admixture from both Neandertal and Denisovans (Prufer et al. 2014; Sankararaman et al. 2014; Sankararaman et al. 2016), we also examined in our dataset the influence of archaic admixture on depressive symptoms, previously investigated in European populations (Simonti et al. 2016).

## Results

The CONVERGE study of Major Depressive Disorder (MDD) previously recruited and sequenced the genomes of 11,670 Han Chinese women at a median coverage of 1.7X (range 0.7-4.3) per individual, of which 10,640 individuals and 25,057,223 SNPs remained after quality control (CONVERGE 2015; Cai et al. 2017). Comparing the allele frequency to East Asian sample in Exome Aggregation Consortium (ExAC; (Lek et al. 2016)), we find extremely high correlation despite potential fin-escale differences in East Asian ancestry (avg correlation coefficient of 0.995 across 22 autosomes and X chromosome). Restricting analysis to variants with minor allele counts (MAC) greater than ten (9,888,655 variants), we find that 548,401 (5.5%), 645,027 (6.5%), and 944,213 (9.5%) are not found in 1000G (Phase 3), 1000G (Phase 3) East Asians, and 1000G (Phase 3) CHB+CHS, respectively [FIGURE 1]. As expected, a large proportion (75-87%) of novel variants were present at low frequency in the population (< 0.05), and 59-75% of them were < 0.005. We also identified a large number of novel variants with high allele frequencies: 13-23% at a frequency of greater than 0.05, which may potentially enable better resolution in genome-wide association studies on common alleles in the Han Chinese population. Figure 1 shows the distribution of the novel variants as a function of allele frequency. **Table S1** gives the number of variants called per chromosome from sequencing data and the percentage of common (>0.05), low frequency (<0.05) and rare (<0.005), known and novel variants passing variant quality score filtering.

**Figure 1:**
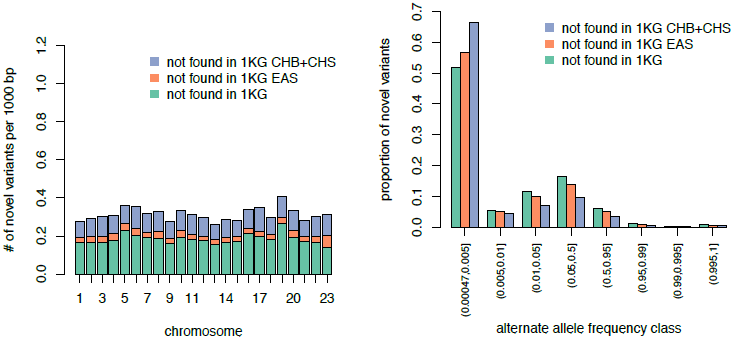
distribution of variant density and frequency spectra of novel variants discovered. Novel variants are defined with respect to three increasing level of inclusiveness: those that are not found among 1KG CHB+CHS populations, 1KG East Asian populations, and all 1KG populations (phase 3). (Left) Density of novel variants discovered per 1000 bp, per chromosome. Chromosome 23 signifies the X chromosome. (Right) Alternate allele frequency of the novel variants.

### Variant function and disease association

We performed variant annotation on the 9.89 million variants with MAC greater than 10 using Variant Effect Predictor (VEP) (McLaren et al. 2016). Over the entire cohort, we found 2,161 Loss of Function (LoF) variants based on VEP (initiator codon variant, splice acceptor variant, splice donor variant, stop lost, and stop gain variants), 12,458 damaging variants predicted by PolyPhen (probably and possibly damaging), and 10,277 damaging variants predicted by SIFT. In total, 17,768 variants are annotated as functionally deleterious by at least one method, of which, 5,633 (31.7%) are common, 4,246 (23.9%) are low frequency and 7,889 (44.4%) are rare [**Figure S1**]. In comparison, we do not see the same degree of enrichment of rare variants among variants with less severe or deleterious effects (missense or synonymous variants in VEP, benign and tolerated variants predicted by PolyPhen and Sift, respectively), consistent with stronger negative selection experienced by the functionally deleterious alleles [**Figure S1**].

At the individual level, we counted the number of alternative alleles (with respect to reference genome) per individual. We find an average of 3,708 functionally deleterious alleles per individual. Most of these alleles are common in the population (91% would be common in the population, 7.6% low-frequency and 1.6% rare), as most of the deleterious load per person is due to the common variants. Similarly, we see that there is a depletion of rare alleles per individual, compared to the less deleterious variant classes such as synonymous variants [**Table S2**].

We investigated further the functional consequences of the variant map that we have generated by cross-referencing the dataset with ClinVar. In total, ClinVar records 56,446 sites reported to be pathogenic or likely pathogenic. ExAC has reported a number of erroneous entries in ClinVar (and other similar database) by showing a number of these variants to be too high in frequency in large aggregate unascertained population of predominantly European ancestry. Of the 56,446 sites in ClinVar, we find that 15,157 sites overlapped ExAC (28.2%). Among these, five are found to be rare in Non-Finnish Europeans in ExAC, but quite frequent in the CONVERGE dataset [Table 1]. This includes rs2276717 in SLC7A14 for retinitis pigmentosa. The frequency of this variant is likely too high in Han Chinese populations to be truly pathogenic, considering the prevalence of the disorder in Han Chinese. We also identified CLINVAR variants that are not covered in ExAC, but are common among the Han Chinese [**TABLE S3**]; the non-exonic nature of these variants present additional challenge in their interpretation, and their pathogenicity should be further scrutinized in the future.

**Table 1:** List of rare pathogenic variants in Europeans but common in CONVERGE. Of the 15,157 sites in CLINVAR reported as Pathogenic or Likely Pathogenic and found in ExAC, we filtered by frequency of <= 0.01 in ExAC Non-Finnish Europeans (NFE) and >= 0.03 in CONVERGE Han Chinese. Frequencies in ExAC East Asians (EAS) are also given for comparisons.

|  |  |  |  |  | ClinVar Annotations |  |  |  | Frequency Information |  |  |  |
| --- | --- | --- | --- | --- | --- | --- | --- | --- | --- | --- | --- | --- |
| Chr | Pos | rsID | Ref | Alt | Variant Type | Affected Gene | Clinical Significance | Syndrome Name | 1000G CEU | ExAC NFE | ExAC EAS | CONVERGE |
| 3 | 170201230 | rs2276717 | C | T | missense | SLC7A14 | Pathogenic | Retinitis pigmentosa 68 | 0 | 1.50E-05 | 0.026 | 0.048 |
| 6 | 46677098 | rs76863441 | C | A | missense | PLA2G7 | Pathogenic | Platelet-activating factor acetylhydrolase deficiency | 0 | 3.50E-04 | 0.064 | 0.063 |
| 12 | 6925407 | rs28919570 | C | T | missense | CD4 | Pathogenic | Okt4 epitope deficiency | 0 | 0.002 | 0.030 | 0.039 |
| 12 | 112241766 | rs671 | G | A | missense | ALDH2 | Pathogenic | Acute alcohol sensitivity | 0 | 6.20E-05 | 0.266 | 0.051 |
| 13 | 20763612 | rs72474224 | C | T | missense | GJB2 | Likely pathogenic | Nonsyndromic hearing loss and deafness | 0 | 0.002 | 0.072 | 0.056 |

### Population Structure of Han Chinese

Compared to previous genetic investigations of the Chinese population, CONVERGE has one of the broadest sampling of Han Chinese across China. In total, CONVERGE has recruited individuals born in all four metropolis municipalities in China (Beijing, Tianjin, Shanghai, and Chongqing) as well as individuals self-reported to be born in 19 out of 22 provinces and 1 (Guangxi) out of 5 autonomous regions in China [FIGURE 2].

**Figure 2:**
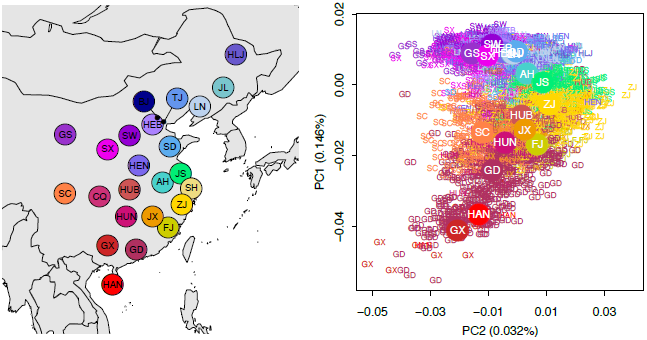
population structure of Han Chinese in CONVERGE. (Left) The geographical locations of each province of China represented in the CONVERGE dataset. The following abbreviations are used: AH, Anhui; BJ, Beijing; CQ, Chongqing; FJ, Fujian; GS, Gansu; GD, Guangdong; GX, Guangxi; HAN, Hainan; HEB, Hebei; HLJ, Heilongjiang; HEN, Henan; HUB, Hubei; HUN, Hunan; JS, Jiangsu; JX, Jiangxi; JL, Jilin; LN, Liaoning; SX, Shaanxi; SD, Shandong; SH, Shanghai; SW, Shanxi; SC, Sichuan; TJ, Tianjin; ZJ, Zhejiang. Beijing, Chongqing, Shanghai, and Tianjin are the four metropolitan municipalities of China; the remaining represent 19 out of the 22 provinces of China. (Right) The first two principal components showing the North-South structure on PC1 and East-West structure on PC2. Individuals are assigned based on their self-reported birth locations and colored according to the map on the left. Individuals from each of the four municipalities were not plotted.

We first conducted a principal components analysis using ∼430,000 common (MAF > 0.10) SNPs for which we have the highest confidence of genotype accuracy (Cai et al. 2017), and a subset of individuals [**METHODS**]. The first two principal components (PCs) recapitulate the geography of China: the first PC corresponds to the North-South axis of China, while the second PC corresponds to the East-West axis of China [FIGURE 2]. As expected, individuals from the major municipalities exhibit greater spread in PC space, reflecting the more diverse origins of the metropolis populations. Nonetheless, individuals from these municipalities are more or less confined to the particular geographical locale [**FIGURE S2**]. Beyond the first two PCs, there is little structure left observable from PCA [**FIGURE S3**].

The strong North-South influence seen in PCA is also recapitulated in pair-wise Fst between provinces. Consistent with the PCA result, Northern Chinese (Heilongjiang, Jilin, Liaoning, Beijing, Tianjin, Hebei, Gansu, Shanxi, Shandong, Shaanxi, Henan) show relatively little genetic differentiation by Fst. In contrast, central Chinese (Jiangsu, Anhui, Shanghai, Hubei, Zhejiang) shows intermediate level differentiation, while further West (Sichuan, Chongqing) and South (Jiangxi, Hunan, Fujian, Guangdong) the differentiation increases [**FIGURE S4**]. For reference, the range of values we observe across CONVERGE provinces are consistent with the Fst between 1000 Genomes CHB and CHS populations at these sites (∼9.9×10^−-4^), as well as those previously reported using array genotypes (Xu et al. 2009; Chen et al. 2016).

### Signature of Admixture with Western Eurasia, Siberia, and neighboring East Asia populations

To examine evidence of admixture, we merged the CONVERGE dataset with the 1000 Genomes dataset and a global reference dataset genotyped on the human origin array data (Patterson et al. 2012). For each Chinese provinces we performed a test for admixture based on allele frequency covariances between CONVERGE sample and the global reference samples serving as potential ancestral populations [**METHOD**, (Patterson et al. 2012)]. In general, we find geographically localized signals of admixture across China [**TABLE S4**], which are corroborated with an allele-sharing statistic comparing CONVERGE provinces of different geographical regions of China to a baseline Chinese population [**METHOD, FIGURE 3**]. Specifically, we find a unique allelic affinity towards Western Eurasia, particularly from Northeastern Europe (*e.g.* Lithuanian, Estonian, etc.) among the Northwestern provinces of China (Gansu, Shaanxi, Shanxi), and a unique affinity towards Siberia among the Northeastern provinces of China (Liaoning, Jilin, and Heilongjiang, and to a lesser extent, Shandong) [FIGURE 3]. In contrast, Southern provinces, such as the southwestern province Sichuan and Southern coastal province Guangdong, tend to show little influence from Siberia or Western Eurasia, but instead tend to have influences from the ethnic minorities geographically situated in the Southwest and Southeast of China, such as the Ami and Atayal from Taiwan, and the Dai [FIGURE 3].

**Figure 3:**
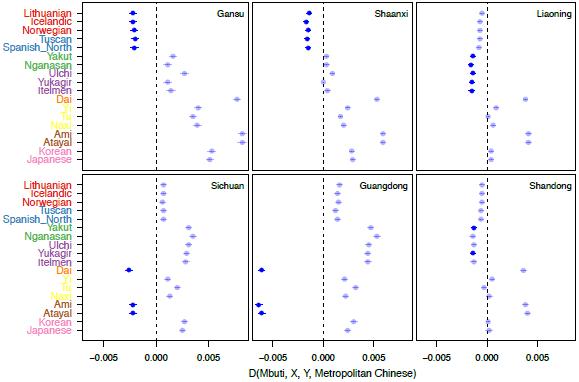
allelic sharing between selected Chinese provinces and non-Han Chinese populations in the Human Origins Array data. The allelic sharing between Han Chinese from Gansu, Shaanxi, Liaoning, Sichuan, Guangdong, and Shandong with various non-Han Chinese populations are evaluated by computing the D-statistics D(Mbuti, X, Y, Metropolitan Chinese), where population X is a population shown on the ordinate, and population Y is one of the Chinese provinces. Populations on the ordinate are color-coded by subcontinental origins: red, Northern Europeans; blue, Southern Europeans; green, Central Siberia; purple, Eastern Siberia; Yellow, Western Chinese minorities; brown, Aboriginal Taiwanese; pink, Korean and Japanese. Metropolitan Chinese is a random sample of 200 individuals from one of the four metropolis municipalities, taken as representative Chinese. A significantly negative value (bolded, Z < −5) suggest significant sharing between populations X and Y, relative to a representative metropolitan Chinese, thus is consistent with a regional signal of admixture.

We also examined whether Han Chinese represented in the CONVERGE sample could be the source of population admixture in neighboring East Asian populations, such as Koreans, Japanese, and Taiwanese [**TABLE S4**]. Koreans have a history of political and demographic relationship with mainland China, and are one of the officially recognized ethnic minority in China, as are Chinese in Korea. Our analyses are consistent with the Koreans receiving genetic influences from China, Siberia, and Japan [**TABLE S4**], but curiously the Chinese provinces, even those in the North that are close to Korean peninsula, did not show signal with Korean as a plausible admixing source, suggesting a largely unidirectional gene flow to Korea. In contrast, we detected no discernable signal of admixture among the island populations of Hainan, Ami and Atayal ethnic minorities from Taiwan, and Japan. This could be due to small sample size, or long-term isolation among the island ethnic minorities that decreases the power of the f3 test for admixture (Patterson et al. 2012). We did, however, identify nine CONVERGE individuals, primarily from the Northeastern provinces, exhibiting affinity for 1000 Genomes JPT population in PCA analysis of merged CONVERGE and 1000 Genome East Asian dataset (Data not shown). These 9 outliers are not otherwise obvious in PCA with CONVERGE alone, but when grouped we find in them evidence for admixture between Japanese and primarily Northern Chinese [**TABLE S4**], consistent with possibly recent admixture from Japan.

### Genetic relationship with ancient human populations

The array of ancient DNA data now publicly available has informed tremendously the genetic ancestry of modern day Europeans. We therefore assessed the genetic relationship between Han Chinese in CONVERGE and 319 publicly available pre-historic humans samples organized into 101 populations (Lazaridis et al. 2014; Fu et al. 2016; Lazaridis et al. 2016). Geographically, the ancient samples spanned three broadly defined geographic regions (Europe/Anatolia, Central Asia, and Siberia) [**METHOD**]. Temporally, the ancient human samples span a period from ∼2,300 years before present to ∼45,000 years before present, with most samples originating since the Mesolithic times ∼9,000 years ago.

We measured allele frequency covariances between ancient populations and CONVERGE provinces using the outgroup f3 statistics (Raghavan et al. 2014). This measure estimates the amount of shared history, or shared drift, between the two populations related through a tree rooted by the outgroup population. In general, we find the shared drift with CONVERGE is greater among Siberian, Central Asian, and Northern European ancient individuals than individuals from Southern Europe, Anatolia, Levant, and the Caucasus [FIGURE 4]. Across CONVERGE, shared drift tends to be significantly correlated with latitude, with Northern provinces showing higher values than Southern provinces [**Table S5**]. This likely reflects the enrichment of European / Siberian ancient samples currently available. The exception is with some of the oldest samples (*e.g.* Ust Ishim, Oase1, Kostenki14 and the archaic individuals), which showed very little variation in the shared drift parameter across the CONVERGE sample, consistent with these oldest samples being a lineage basal to the European-East Asian pre-divergence lineage, while the more recent samples showed greater variability across geography [**FIGURE S5**].

**Figure 4:**
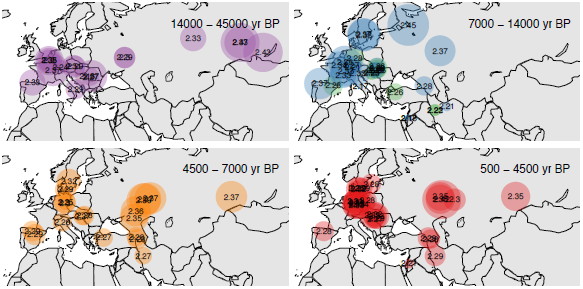
shared drift between Han Chinese and ancient DNA samples from Eurasia. Using Shanghai individuals as representatives, shared drift between Chinese and ancient humans arecomputed by calculating the outgroup f3 statistics of the form f3(Mbuti; X, Y). Ancient individuals are separated into roughly four temporal categories spanning from 500 years before present to 45,000 years before present. In the time period 7,000 to 14,000 years before present, ancient individuals are further divided based on their cultural context: pre-Neolithic hunter---gatherers in,blue, Neolithic farmers in green. During this period we find modern Chinese individuals showing greater shared drift with pre-Neolithic hunter-gatherers rather than Neolithic farmers.

Spatiotemporally, before the major warming period (∼14000 years BP), some of the largest shared drift values observed were between the three Siberian ancient individuals (MA1, AG2, AG3, shared drift = 0.241, 0.240, and 0.236, respectively) and CONVERGE [FIGURE 4]. During the transition period from Paleolithic/Mesolithic to early Neolithic where a mixture of both hunter-gatherers and Neolithic farmers are found in Europe (∼7000-14000 years BP), the individuals showing the greatest shared drift with CONVERGE samples are the pre-Neolithic hunter-gatherer individuals (blue), rather than the Neolithic individuals (green) [FIGURE 4]. In particular, the Karelia hunter-gatherers, Motala hunter-gatherers, and the Loschbour hunter-gatherer across the Eastern, Northern, and Western Europe had the highest values of shared drift during this time period (outgroup f3 statistics = 0.243, 0.236, and 0.236, respectively; standard errors = 0.0029, 0.0028, and 0.0031, respectively), compared to Neolithic individuals (outgroup f3 statistics of 0.216 to 0.228; standard errors of 0.002 to 0.003). The more intermediate values observed for later time periods could then reflect the documented mixing between huntergatherers, Eastern pastoralists, and Neolithic farmers in Europe and their subsequent dispersal (Lazaridis et al. 2014; Haak et al. 2015; Mathieson et al. 2015). Together, these observations suggest that the arrival of farming in China followed a different pattern/mechanism as it did in Europe.

### Genetic relationship with archaic hominin individuals

Past studies of Neandertal genomes showed the East Asians have inherited ~20% more Neandertal ancestry than Europeans have and that this excess ancestry may reflect a second pulse of admixture in East Asians or a dilution of Neandertal admixture in Europeans (Prufer et al. 2014; Sankararaman et al. 2014; Vernot and Akey 2014; Kim and Lohmueller 2015; Vernot and Akey 2015; Sankararaman et al. 2016). We largely recapitulated the relationship of a number of Neandertal samples and Denisovan to the Han Chinese as previously reported [**FIGURE S6**]. We observed subtle allele-sharing and estimated Neandertal ancestry (approximately 1.8% to 2%) differences across China, though the difference is not significant after correcting for multiple testing [**TABLE S6**].

Previous analyses of the locations of Neanderthal segments within the genomes of non-African individuals indicate that some of the Neanderthal variants were adaptively beneficial while the bulk of Neanderthal variants were deleterious in the modern human genetic background (Harris and Nielsen 2016; Juric et al. 2016). Specifically, a recent examination of Neandertal-informative markers (NIMs) among large cohort of Europeans showed that these markers explained some proportions of the phenotypic risk of a number of diseases from electronic health record (Simonti et al. 2016), including MDD. We sought to replicate that finding in East Asians.

We extracted 75,539 SNPs that were previously identified to tag Neandertal haplotypes in East Asian individuals in Phase 1 of the 1000 Genomes project (Sankararaman et al. 2014), and assessed the contribution of these NIMs to depression in the CONVERGE cohort consisting 5,224 cases of MDD and 5,218 controls. The allele frequencies of these NIMs are highly correlated (r = 0.951) between CONVERGE sample and 1000 Genomes, suggesting that the NIMs tested here are not impacted by the low coverage sequencing. We tested the association between the NIMs and depression by performing a logistic regression of depression, controlling for age and the first ten principal components, for each of the two traits. We found no association surviving the Bonferroni correction [**FIGURE S7**] and the QQ plots did not reveal any systematic inflation nor significant enrichment among top associated SNPs [data not shown].

We also calculated the proportion of phenotypic variance explained by these NIMs using GCTA (Yang et al. 2011) for MDD. We used a prevalence of 7.5% for MDD to transform the heritability to the liability scale. We find that the variance explained by the NIMs is ∼1%, which is different from that reported in Simonti et al. (∼2%) and is not significantly different from 0 (P = 0.12). Repeating the analysis with NIMs with MAF > 0.01 as well as with no covariates did not qualitatively alter the results [**TABLE S7**]. Finally, we evaluated whether the contribution of the NIMs to the heritability of MDD is more or less than expected based on the genetic architecture of MDD. We found that the heritability explained by NIMs is not significantly different from that of a background set of SNPs chosen at random to match the NIMs by derived allele frequency decile and by LD scores (P > 0.4).

### PCA-BASED SIGNAL OF SELECTION

Taking advantage of the greater geographical resolution of CONVERGE, we conducted a genome scan for loci under directional selection within the Chinese populations. We used a recently published PCA-based method that identifies the most differentiated SNPs along each PCs (Galinsky et al. 2016). We identified 24 loci showing genome-wide significant allele frequency differentiation across CONVERGE sample in the top 2 PCs [**Figure S8**]. Some of the loci appeared significant in both PCs [Table 2, Table 3], and a number of them had been observed to lie in the tail end of haplotype-based selection statistics in East Asian populations [Table 2, Table 3], thereby confirming the robustness of our approach here. Comparing pattern of uniquely shared sites with archaic humans (Neandertal from Altai, Vindija, and Denisovan) at these loci and the rest of the genome suggests these loci are not due to archaic introgression [**Methods**], consistent with their being selected after the diversification of Han Chinese. In particular, a number of our top loci are known to be related to diet, UV radiation, and immune responses, the major selective forces known to impact global human populations. However, previous studies are often based in non-East Asian populations, or compared East Asian populations as a whole to non-East Asian populations; our results here suggest these loci show within-China signal of selection.

**Table 2:** putative loci under selection on PC1. Locus intervals are defined as the interval at which the P-value for evidence of selection drops below 0.05. Lead SNP is the SNP with the best P-value after fine-mapping, which included all SNPs with MAF >= 1% within each interval. If the interval overlapped with previous selection scans involving East Asian samples, the references are given (Hirayasu et al. 2008; Suo et al. 2012; Liu et al. 2013); ^*^ denote corroboration by a haplotype-based statistic. Notable genes column lists only protein coding genes for which a variant with P < 1e-7 was mapped and annotated by VEP to its canonical transcript. Notable phenotype associations are phenotypes reported to be associated with the genes in the GWAS catalog (Welter et al. 2014).

| Chr | Start (Mb) | Stop (Mb) | Locus Size (Mb) | Lead SNP | P-value | Previously reported? | Notable Genes | Notable Phenotype Associations |
| --- | --- | --- | --- | --- | --- | --- | --- | --- |
| 1 | 11.84 | 12.00 | 0.161 | 1:11856378 | 4.45E-11 |  | C1orf167, MTHFR, CLCN6 | Homocysteine levels |
| 1 | 207.66 | 207.80 | 0.143 | 1:207694357 | 2.07E-08 |  | CR1 |  |
| 3 | 75.59 | 75.76 | 0.163 | 3:75632610 | 9.59E-12 |  |  |  |
| 3 | 162.37 | 163.25 | 0.881 | 3:162732731 | 1.41E-14 |  |  |  |
| 6 | 29.72 | 30.47 | 0.747 | 6:29878687 | 5.26E-13 | MHC, Liu et al.*, Suo et al.* | MHC region |  |
| 6 | 31.31 | 31.35 | 0.04 | 6:31313972 | 1.53E-10 | MHC, Liu et al.*, Suo et al.* | MHC region |  |
| 6 | 32.68 | 32.68 | 0.001 | 6:32682207 | 1.16E-08 | MHC, Liu et al.*, Suo et al.* | MHC region |  |
| 11 | 61.52 | 61.69 | 0.176 | 11:61579463 | 7.80E-14 | Suo et al.* | MYRF, TMEM258, FEN1, FADS1, FADS2, FADS3, RAB3IL1 |  |
| 13 | 99.60 | 99.77 | 0.17 | 13:99759875 | 2.02E-11 | Suo et al.* | DOCK9, DOCK9-AS2 |  |
| 14 | 105.91 | 106.38 | 0.479 | 14:106134635 | 9.34E-32 |  | IGH cluster |  |
| 15 | 24.34 | 24.97 | 0.639 | 15:24341809 | 1.26E-22 |  | PWRN2, PWRN3, PWRN1, NPAP1 |  |
| 19 | 54.74 | 54.80 | 0.057 | 19:54800371 | 2.80E-26 | Hirayasu et al. | LILRB5, LILRB2, LILRA3 |  |
| 20 | 0.77 | 0.78 | 0.011 | 20:773680 | 1.09E-08 |  |  |  |

**Table 3:**
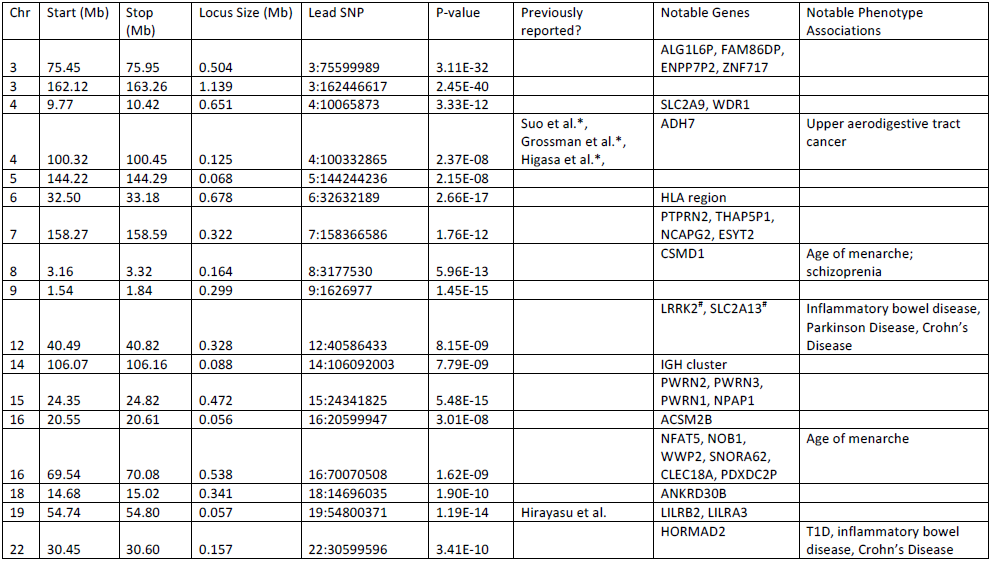
putative loci under selection on PC2. Locus intervals are defined as the interval at which the P-value for evidence of selection drops below 0.05. Lead SNP is the SNP with the best P-value after fine-mapping, which included all SNPs with MAF >= 1% within each interval. If the interval overlapped with previous selection scans involving East Asian samples, the references are given (Hirayasu et al. 2008; Higasa et al. 2009; Suo et al. 2012; Grossman et al. 2013); ^*^ denote corroboration by a haplotype-based statistic. Notable genes column lists only protein coding genes for which a variant with P < 1e-7 was mapped and annotated by VEP to its canonical transcript. Notable phenotype associations are phenotypes reported to be associated with the genes in the GWAS catalog (Welter et al. 2014); ^#^ denotes cases where no genes were identified based on the above criteria, and thus the closest genes are given.

| Chr | Start (Mb) | Stop (Mb) | Locus Size (Mb) | Lead SNP | P-value | Previously reported? | Notable Genes | Notable Phenotype Associations |
| --- | --- | --- | --- | --- | --- | --- | --- | --- |
| 3 | 75.45 | 75.95 | 0.504 | 3:75599989 | 3.11E-32 |  | ALG1L6P, FAM86DP, ENPP7P2, ZNF717 |  |
| 3 | 162.12 | 163.26 | 1.139 | 3:162446617 | 2.45E-40 |  |  |  |
| 4 | 9.77 | 10.42 | 0.651 | 4:10065873 | 3.33E-12 |  | SLC2A9, WDR1 |  |
| 4 | 100.32 | 100.45 | 0.125 | 4:100332865 | 2.37E-08 | Suo et al.*, Grossman et al.*, Higasa et al.* | ADH7 | Upper aerodigestive tract cancer |
| 5 | 144.22 | 144.29 | 0.068 | 5:144244236 | 2.15E-08 |  |  |  |
| 6 | 32.50 | 33.18 | 0.678 | 6:32632189 | 2.66E-17 |  | HLA region |  |
| 7 | 158.27 | 158.59 | 0.322 | 7:158366586 | 1.76E-12 |  | PTPRN2, THAP5P1, NCAPG2, ESYT2 |  |
| 8 | 3.16 | 3.32 | 0.164 | 8:3177530 | 5.96E-13 |  | CSMD1 | Age of menarche; schizophrenia |
| 9 | 1.54 | 1.84 | 0.299 | 9:1626977 | 1.45E-15 |  |  |  |
| 12 | 40.49 | 40.82 | 0.328 | 12:40586433 | 8.15E-09 |  | LRRK2<sup>#</sup>, SLC2A13<sup>#</sup> | Inflammatory bowel disease, Parkinson Disease, Crohn's Disease |
| 14 | 106.07 | 106.16 | 0.088 | 14:106092003 | 7.79E-09 |  | IGH cluster |  |
| 15 | 24.35 | 24.82 | 0.472 | 15:24341825 | 5.48E-15 |  | PWRN2, PWRN3, PWRN1, NPAP1 |  |
| 16 | 20.55 | 20.61 | 0.056 | 16:20599947 | 3.01E-08 |  | ACSM2B |  |
| 16 | 69.54 | 70.08 | 0.538 | 16:70070508 | 1.62E-09 |  | NFAT5, NOB1, WWP2, SNORA62, CLEC18A, PDXDC2P | Age of menarche |
| 18 | 14.68 | 15.02 | 0.341 | 18:14696035 | 1.90E-10 |  | ANKRD30B |  |
| 19 | 54.74 | 54.80 | 0.057 | 19:54800371 | 1.19E-14 | Hirayasu et al. | LILRB2, LILRA3 |  |
| 22 | 30.45 | 30.60 | 0.157 | 22:30599596 | 3.41E-10 |  | HORMAD2 | T1D, inflammatory bowel disease, Crohn's Disease |

A well-known selective pressure in human evolution is human interaction with pathogens. We identify a number of signals related to immune response. In addition to the expected signal from the HLA region, three known selected loci reached genome-wide significance in our scan. These include the *LILRA3* / *LILRB2* locus on chr 19, which encodes a family of HLA class-I recognizing receptors, the *CR1* locus on chr 1, which encodes the human complement receptor 1 gene, and the IGH locus on chr 14, the Immunoglobulin Heavy cluster [**FIGURE S9-S11**] (Cockburn et al. 2004; Barreiro et al. 2008; Hirayasu et al. 2008; Galinsky et al. 2016). The latter two signals appear to be due to a variant different from previously reported signal in other populations. Adaptation to different diets is another known selective force of human evolution. We find two loci related to diet reaching genome-wide significance in our scan: the *FADS1,2,3* locus on chr 11 and the *ADH* gene cluster on chr 4. The FADS locus encodes a cluster of fatty acid desaturases which modulate omega-3 polyunsaturated fatty acids levels, and is known to be selected in Europeans and Inuits (Fumagalli et al. 2015; Mathieson et al. 2015). The *ADH* locus encodes a cluster of alcohol dehydrogenases involved in alcohol metabolism. However, our lead SNP resides in *ADH7*, is in poor LD (r2 ∼ 0.18) with the previously identified variant in *ADH1B* that were shown to be under selection in East Asians and Europeans (Han et al. 2007; Li et al. 2011a; Galinsky et al. 2016), and thus may suggest a different Han-Chinese specific signal of selection. Lastly, we note a signal on chr1 in the *MTHFR* gene encoding a methylenetetrahydrofolate reductase involved in folate metabolishm [TABLE 2]. This signal may represent a complex interaction between UV radiation (to prevent photolysis of folate) and dietary folate intake (Rosenberg et al. 2002; Jablonski and Chaplin 2010).

## Discussion

By analyzing the low-coverage whole-genome sequences of 11,670 Han Chinese individuals, we produced and characterized currently the largest and most comprehensive map of genetic variation in the Han Chinese population. We have used this catalog to investigate population structure, admixture history, and infer signals of natural selection. Our results add to the existing knowledge of this previously relatively understudied, yet one of the largest, population in the world, and pave way to future medical and population genetic studies of Han Chinese.

Analyses of exome sequencing studies, primarily in Europeans, have noted a number of erroneous entries in clinical genetics databases, such as ClinVar, where a reported pathogenic variant appears to be too common in the population to be truly pathogenic. We cross-referenced the catalog of Han Chinese variation with the ClinVar database, looking for incidences where reported pathogenic variants are extremely rare in European populations (hence likely to be still considered pathogenic based on ExAC), but are more common in Han Chinese. We identified 5 pathogenic or likely pathogenic variants in the ClinVar database where the estimated frequency in ExAC non-Finnish Europeans is < 0.01, but the estimated frequency in Han Chinese is > 0.03 [TABLE 1], and one additional likely pathogenic allele for a rare Mendelian disease in the intronic region that was not covered in ExAC but is common in Han Chinese [**TABLE S2**]. Among them, the missense variant in SLC7A14 (rs2276717) is a singleton in ExAC NFE (MAF ∼1.5x10^−5^), but has a frequency of 0.048 in Han Chinese. The variant is reported to cause an autosomal recessive form of retinitis pigmentosa (Jin et al. 2014). However, given the prevalence of approximately 1 in 4000 in China (You et al. 2013), which is on par with the estimates in U.S. (<u>https://nei.nih.gov/health/pigmentosa/pigmentosa_facts</u>), the maximum credible population allele frequency would be approximately 0.022 in Han Chinese even under relatively relaxed assumptions of genetic architecture [**METHOD**]. Therefore, this variant is too common and unlikely to be pathogenic in the Han Chinese population. Incidentally, the observed frequency in ExAC East Asians is lower (0.027, but likely contains a mixture of Han Chinese and other East Asian populations), and much closer to the maximum credible population allele frequency, and thus would not be as confidently filtered in a screen for pathogenic variants. Similarly, rs11085825 in the intron of GCDH gene is likely too common in Han Chinese (and in fact, across the world based on 1000 Genomes) to be responsible for the autosomal recessive disorder of glutaric aciduria, type 1 (occurs in ∼1 per 100,000 infants (Hedlund et al. 2006)), despite being reported likely pathogenic in ClinVar.

Despite these examples where examining the Han Chinese populations helps filter out candidate pathogenic variants, interpretation of the other entries in ClinVar is less obvious. For example, rs671 in ALDH2 for alcohol dependence, is a well-characterized functional variant in the alcohol metabolism pathway, has been suggested to be selected in East Asian populations (Oota et al. 2004), and thus would not be surprising that the frequency is uniquely high in East Asians while being truly causal. For another example, rs72474224, a missense variant in GJB2, is reported as likely pathogenic for autosomal recessive form of non-syndromic sensorineural hearing loss and deafness by ClinVar. The variant is rare in non-East Asian populations in ExAC (< 0.001), but is common in ExAC East Asian (0.072) and CONVERGE Han Chinese (0.055). However, the prevalence of non-syndromic hearing loss is also known to be higher in East Asian populations (Naeem and Newton 1996), and thus the observed frequency in Han Chinese may still be consistent with the its pathogenicity. Taken together, these findings suggest that population-specific prevalence and disease risk should be taken into account when screening clinically sequenced genomes for pathogenic variants.

We also utilized this whole-genome sequencing resource to address a number of basic population genetic questions in this population. Specifically, we examined population structure, signals of admixture, relationships with ancient and archaic humans, and signals of allelic differentiation consistent with positive selection. Because of the ultra-low coverage in generating this dataset, we focused on using allele-frequency based methods to address these questions.

The North-to-South structure among the Han Chinese has been a well-known feature, evident from both studies of uniparental markers (Wen et al. 2004; Zhao et al. 2015) and genome-wide chip array data (Chen et al. 2009; Xu et al. 2009; Chen et al. 2016). However, little structure have been observed beyond the first principal component, which could be due to a combination of smaller sample sizes in the previous studies, as well as the lack of breath in representing present day China. We observed, for the first time to our knowledge, clear population structure in the East-to-West axis among the CONVERGE sample. The E--W pattern is clearly weaker, explaining only ∼1/5 of the variance explained by the first principal components. However, this implies that while there may be increased migration across longitude in the spread of the Han Chinese, the migration pattern follows sufficiently an isolation-by-distance model to be detectable in a large well-represented sample.

We applied a test of admixture based on the f3 statistics (Patterson et al. 2012), which utilizes allele frequency covariances between CONVERGE sample and our chosen global reference samples (the Human Origins Array data) to test for evidence of admixture. We find patterns of admixture signals that are geographically localized to certain parts of China [**TABLE S4**]. In general, the admixture signals that we detected are also corroborated by allelic-sharing statistics [Figure 3], and make sense geographically in that the signals tend to be with neighboring non-Han Chinese populations that are situated closely. The signal we have detected through f3-statistics is thus likely again demonstrating a pattern of isolation-by-distance, with constant / frequent exchange of migrants with neighboring populations through history, rather than a single pulse admixture model typically invoked for populations such as the African Americans. This suggests that caution is needed to interpret the admixture results via the f3 statistics (Peter 2016).

One finding from our analysis of admixture signals that most likely fit a one-pulse admixture model is our observation of admixture from Northern European populations to the Northwestern provinces of China (Gansu, Shaanxi, Shanxi), but not other parts of China. Previous analysis of the HGDP data, based on patterns of haplotype sharing among 10 Han Chinese from Northern China, estimated a single pulse of ∼6% West Eurasian ancestry among the Northern Han Chinese. The estimated date of admixture was around 1200 CE. This signal is also observed among the Tu people, an ethnic minority also from Northwestern China; the authors attributed this signal to contact through the Silk Road (Hellenthal et al. 2014). We estimate a lower bound of admixture proportion due to Northern Europeans at approximately 2% - 5%, with an admixture date of about 26 +/-3 generations for Gansu, and 47 +/-3 generations for Shaanxi [**Table S8**]. Using a generation time of about 26-30 years (Moorjani et al. 2016), these estimates correspond to admixture events occurring at around 700 CE and 1300 CE, respectively, corresponding roughly to the Tang and Yuan dynasty in China. However, these estimated dates should be interpreted with caution, as both the violation of a single pulse admixture model and the additional noise in inter-marker LD estimates due to low coverage data could bias the estimates.

The prevailing view in the origin of agriculture in China is that domestication of crops and animals occurred in China independent of other agricultural centers around the world, such as the Fertile Crescent (Ho 1976; Jones and Liu 2009; Zhao 2011). Consistent with this view, and unlike how agriculture arrived in Europe (due to demic diffusion from Anatolia (Mathieson et al. 2015)) and South Asia (due to demic diffusion from Iran (Lazaridis et al. 2016)), we find that present day Han Chinese show greater genetic affinity to Mesolithic European hunter--gatherers than Neolithic European farmers from the same era [FIGURE 4], and that they show very low affinity with the Iranian Farmers [FIGURE 4]. Currently available ancient DNA data are largely of European origin, thus have limited information on the spread of the Han Chinese people that appears to be coupled with spread of agriculture (Zhao et al. 2015). Furthermore, within China there may be two or more independent centers of domestications: one near the southern Yangtze River where rice was first domesticated, and another one near the northern Yellow River where millets are found (Zhao 2011). Therefore, the extent to which the Han Chinese people replaced or were assimilated during their expansion and how that coupled with the spread of agricultural practices may be of interest to study in the future with more ancient DNA data being generated out of China.

Taking advantage of the most geographically diverse dataset of Han Chinese, we identified a number of loci in the genome showing extreme frequency differentiation across China, a subset of which is likely due to selective pressure such as pathogen, diets, and variation in environmental exposures such as UV radiation. A few of these loci have been well studied, and are highlighted below.

The FADS locus was recently identified to be under selection in the Greenlandic Inuits, presumably driven by the marine--rich diets of the arctic population (Fumagalli et al. 2015), as well as in Europeans (Mathieson et al. 2015). Our top variants in the FADS locus, two intronic variants in FADS1 (chr11:61571478 and chr11:61579463, r2 = 0.99 between the two), are in high LD in CONVERGE cohort with one of the top hits reported in Fumagalli et al. (chr11:61597212; r2 = 0.99), while the other five variants reported in Fumagalli et al. showed modest to poor LD with our lead variant here [**Figure S12, Table S9**]. This may reflect a common demographic origin between the two peoples (perhaps mediated through common ancestry prior to the Inuit’s expansion across Beringia). However, because the presumed selected (derived) allele in Inuits for chr11:61597212 is actually in higher frequency in Southern Han Chinese rather than Northern Chinese, it remains possible that this is an instance of convergent selection on standing variation due to shared marine diet of the Inuits and Southern Han Chinese. Alternatively, it could also be due to selection of the alternative allele in Northern Chinese, as there is evidence of varying selection on the two alleles in FADS locus depending on different dietary practices in Europe (Ye et al. 2017). Future analysis focusing on the haplotypic pattern of selection will provide more insight to the allelic relationship observed in Han Chinese and Inuits at this locus.

Another well-studied locus that our selection scan detected is in MTHFR. Our top differentiated SNP is a missense variant (rs1801133, 1:11856378) that has been shown to modulate the activity of the enzyme by producing a thermolabile derivative with reduced activity (Frosst et al. 1995) and thus is associated with plasma homocysteine levels (Pare et al. 2009; van Meurs et al. 2013). The thermolabile allele frequency ranges from ∼30% to ∼60% in CONVERGE samples from Southern and Northern China, respectively, consistent with previously reported South-to-North increasing gradient in East Asia and in opposite direction to the observed North-to-South increasing gradient in Europe (Yafei et al. 2012). At high latitude, less pigmented skin is selectively favored to improve vitamin D production, but would in turn be susceptible to photolysis of folate. Therefore, the non-thermolabile alleles would be favored to more efficiently metabolize folic acid (Jablonski and Chaplin 2010), leading to the North-to-South increasing gradient of the thermolabile allele. The trend is opposite in East Asia, but could still be explained by the relative difference in latitude between China and Europe (Yafei et al. 2012). Furthermore, dietary supplement of folate could stabilize the thermolabile protein (Frosst et al. 1995), and in cases where folate is plentiful in the diet the impact of the thermolabile allele on recurrent pregnancy loss is reduced (Munoz-Moran et al. 1998). Thus, differential dietary practices across Han-Chinese could lead to the latitudinal gradient we observe here. However, UV exposure appears to better explain the allele frequency gradients than dietary practices in China (Yafei et al. 2012).

We also detected a signal at the ADH locus, which is near but different from the well-characterized variant in ADH1B locus (rs1229984, 4:100239319, Arg48His allele) for alcohol metabolism. The ADH1B Arg48His variant is suggested to be positively selected based on comparisons between East Asians and Europeans (Han et al. 2007), as well as within Europeans (Galinsky et al. 2016). Our lead variant (rs422143, 4:100332865) is downstream of *ADH7*, approximately 90kb away from *ADH1B*, and showed much stronger signal than any variant in the *ADH1B* locus (P ∼ 2.4e-8, TABLE P6). It is in poor LD (r2 ∼ 0.18) with the *ADH1B* Arg47His variant, which only showed marginal evidence of association among CONVERGE samples (*P ∼* 0.00096). However, it should be noted that a regulatory variant in the *ADH1B* locus, rs3811801, was suggested to be a younger variant showing stronger differentiation within East Asia than Arg48His variant (Li et al. 2011a; Galinsky et al. 2016); this variant is in slightly higher LD to our lead variant in *ADH7* [r2 ∼ 0.33], as well as better evidence of selection [*P* ∼ 6.1 × 10^−6^].

*ADH7* is a class IV alcohol dehydrogenase; unlike the class I enzymes (*ADH1A, ADH1B, ADH1C*) that are mainly expressed in the liver and account for ∼80% of the post-absorptive alcohol metabolism, *ADH7* is mainly expressed in the upper digestive tract where it oxidizes ethanol at high concentrations early in the timeline of alcohol metabolism (Park et al. 2013). The mechanistic differences could also explain why the associations of variants in *ADH1B* with UADT cancer (squamous cell carcinoma of the upper aerodigestive tract, encompassing oral cavity, pharynx, larynx, and esophagus) is mediated through alcohol consumption behaviors, while the associations of variants in *ADH7* with UADT appears to be independent from alcohol consumptions (McKay et al. 2011). Taken together, these observations are consistent with the *ADH7* signal in our selection scan being independent from the previously reported *ADH1B* signal, and specific to Han Chinese. The potential selective pressures for alcohol dehydrogenase variants in East Asia include protection against infectious agents (due to increased acetaldehyde levels (Han et al. 2007)) and rice domestication (Peng et al. 2010). The latter hypothesis is based on the correlation between ages of archeological sites of rice domestication and allele frequencies of ADH1B Arg48His allele, which spread along the E-W axis of China. Consistent with these observations, our detected signal in ADH7 is found only on PC2, which corresponds to E-W axis. Furthermore, we find enrichment of selection signals in PC2 for another well-characterized locus involved in alcohol metabolism (Brooks et al. 2009), the ALDH2 locus on chr12 (a number of SNPs with P-values ranging from 1×10^−4^ to 1×10^−6^; rs671, 12:112241766, ALDH2 Glu487Lys allele, *P* ∼ 0.00049).

Notably, *ADH7* and *MTHFR* have been recently implicated as under selection in comparison of Han Chinese to Tibetans (Yang et al. 2017). Our results here suggest a more general signal among Han Chinese that may reflect a continuous variation of selective pressure among Han Chinese across geography, rather than a selective pressure specific to the Tibetans. Beyond the loci described here, there are a number of potentially novel loci found with little recognizable insights to the biological mechanisms or the selective pressure behind the extreme differentiation. Some of the genes within the selection peak or nearby have been implicated in GWAS for life history traits such as age of menarche, or immune-related traits such as inflammatory bowel disease or Crohn’s disease, consistent with a signal of selection at these loci, but we note that there may be Han Chinese-specific but yet uncharacterized regions of long range LD that could appear like a region under selection in this analysis.

We have characterized extensively the pattern of population structure, demographic history, and extreme allelic differentiation within a broad and geographically diverse dataset of Han Chinese. However, a major limitation to our study is the reliance on allele frequency based methods for analysis, as the ultra-low coverage data generation obscured the haplotypic patterns at the individual level and precludes direct merging and comparison to other genotyped reference non-Chinese datasets. Therefore, as the cost of sequencing continues to decrease, much of the conclusions drawn here regarding the population history of Han Chinese should be complemented with haplotype-based analysis. For instance, while we revealed East-to-West structure among the Han Chinese, the signal is relatively weak and very little structure is discernable beyond the second PC. Haplotype-based clustering of individuals would produce a finer view of the structure among the Han Chinese (Lawson et al. 2012). The unit of much of our downstream analyses was also based on self-reported birthplaces among the CONVERGE participants. Using Haplotype-based clustering to define the units of analysis among the CONVERGE sample may further improve the power to detect signals of admixture, and enable reliable estimations of admixture proportions and dates. Nevertheless, our results here collectively demonstrate that there exist significant variations in demographic and adaptive histories across Han Chinese populations. Here we demonstrated how the impact on one type of trait, that of MDD and Melancholia, due to Neandertal ancestry appears to differ between the Han Chinese and Europeans. In general, these unique histories undoubtedly contributed to the variation of phenotype within the Han Chinese and between Han Chinese and other global populations. Therefore an better understanding of the Han Chinese history will help conduct and interpret medical genetic studies in the future in this largest ethnic group of mankind.

## Methods

### Sample collection, DNA sequencing and variant calling

CONVERGE collected cases of recurrent major depression from 58 provincial mental health centers and psychiatric departments of general medical hospitals in 45 cities of China. Controls were recruited from patients undergoing minor surgical procedures at general hospitals (37%) or from local community centers (63%). All subjects were self-reported Han Chinese women born in China with four Han Chinese grandparents. The study protocol was approved centrally by the Ethical Review Board of Oxford University (Oxford Tropical Research Ethics Committee) and the ethics committees of all participating hospitals in China. All participants provided their written informed consent. The descriptions of DNA sequencing and variant calling pipelines and access to the sequencing and called genotype data have been previously published (CONVERGE 2015; Cai et al. 2017).

### Annotation with VEP

Called variants across the 22 autosomes and X chromosome were functionally annotated using the variant effect predictor (version 84) in offline mode, with options for SIFT and Polyphen prediction terms for coding variants turned on. Variants with multiple annotations due to overlaps to multiple genomic features are resolved by assigning the most severe annotation based on the order suggested here: <u>http://uswest.ensembl.org/info/genome/variation/predicted_data.html#consequences</u>. We defined loss of function variants by pooling variants with the following consequence classes: initiator_codon_variant, splice_acceptor_variant, splice_donor_variant, stop_lost and stop_gained. We further supplemented this list of variants with those that are predicted to be “deleterious” by SIFT or “probably damaging” and “possibly damaging” by Polyphen; together we refer to this list of variants as the functionally deleterious variants Called variants across the 22 autosomes and X chromosome were functionally annotated using the variant effect predictor (version 84) in offline mode, with options for SIFT and Polyphen prediction terms for coding variants turned on. Variants with multiple annotations due to overlaps to multiple genomic features are resolved by assigning the most severe annotation based on the order suggested.

### Evaluating ClinVar variants

Curated ClinVar database was downloaded from <u>https://github.com/macarthur--lab/clinvar</u> on 04/27/2017. We examined variants annotated as pathogenic or likely pathogenic, with no conflicted reports, with low frequency (< 0.01) in ExAC Non-Finnish Europeans (NFE), but common (> 0.03) in CONVERGE Han Chinese. We also cataloged pathogenic and likely pathogenic variants in ClinVar (n= 56,446) that are not covered in ExAC, but are common (> 0.03) in CONVERGE Han Chinese. Because of its near-absence in non-East Asian populations in ExAC, we used an online application (<u>http://cardiodb.org/allelefrequencyapp/</u>, (Whiffin et al. 2017)) to specifically follow up rs2276717 reported in ClinVar for causing the autosomal recessive retinitis pigmentosa. We assumed a autosomal recessive model, with prevalence of approximately 1 in 4000, genetic heterogeneity of 0.02 (Jin et al. 2014), and a range of possible parameters for the allelic heterogeneity and the penetrance. We find the maximum credible population allele frequency is 0.0224.

### Individual-level filtering for population genetics analysis

As studies of population structure and demography typically do not require the sample sizes necessary for a successful GWAS, we set out to identify a set of individuals with the highest quality genotypes suitable for investigating population genetic parameters. Exploratory PCA on the entire cohort showed a number of individuals clustered on PC2 due to extremely low coverage and, consequently, comparatively poorer imputation qualities. Therefore, we trained a classifier to differentiate individuals that clustered with the extreme outliers on PC2. We trained a classifier with seven QC features on 500 random samples of apparent poor quality (outliers on PC2) and 2000 random samples of apparent good quality (non-outliers). We selected a threshold that would achieve 90% sensitivity of selecting poor quality sample in ten-fold cross-validation. We then used this model to classify the remaining 8,140 CONVERGE individuals not used in the training set. In total, we removed 3,183 individuals (retained 7,457 individuals) for the population genetics analysis through the following filters (with overlapping individuals across filters): 2,764 individuals by the classifier above (including 500 training individuals), 18 individuals with inconsistent or missing demographic labels, 227 individuals with missing sequencing quality measures, 71 individuals due to extended relatedness (removing up to and including 3rd degree relationships by KING v1.4 (Manichaikul et al. 2010)), and 138 population outliers in a second round of PCA analysis (+/-4 s.d. in any of the top 10 PCs).

### Datasets used for population genetic analysis

For investigating population structure within the CONVERGE cohort, we generated a dataset of 7,457 individuals with genotypes at 429,935 SNPs, representing a random selection of 10% of all SNPs in CONVERGE with MAF >= 10%, as the common SNPs have much higher genotyping accuracy in a ultra-low coverage sequencing design. For analyses investigating the genetic relationship between Han Chinese from CONVERGE and reference populations, including ancient and archaic individuals, we merged CONVERGE with the Human Origins Array data (N = 2,583, (Lazaridis et al. 2016)) and the 1000 Genomes phase 3 data (N = 2,504, (1000 Genomes Project et al. 2015)). We further supplemented this merged dataset with ancient samples not found in the public release of Human Origins Array, including ancient Europeans from the Ice Age (Fu et al. 2016; Lazaridis et al. 2016), and the recently released high-coverage Vindija 33.19 individual (http://cdna.eva.mpg.de/neandertal/Vindija/VCF/Vindija33.19/). Dataset merges were done by removing apparent triallelic SNPs, SNPs with inconsistent alleles, and A/T or C/G transversion SNPs, and any SNPs that are not available in any of the datasets. In total, we analyzed 449,336 SNPs shared between Human Origins Array data and CONVERGE, out of the 592,146 SNPs available on Human Origins Array data, across 12,608 individuals.

### Population structure and admixture

Within-CONVERGE principal components analysis (PCA) was conducted using all 7,457 unrelated individuals, each labeled according to their self-reported birthplace at the province level. PCA was performed using EIGENSTRAT (v6.1) after removing one SNP of each pair of SNPs with r2 >= 0.8 (in windows of 50 SNPs and steps of 5 SNPs) as well as SNPs in regions known to exhibit extended long-range LD (Price et al. 2008). In total, 194,398 SNPs were used in PCA.

Weir and Cockerham’s unbiased estimator of Fst was also calculated in pairwise fashion between populations grouped at the province level in CONVERGE, across 37,670 SNPs after removing one SNP of each pair of SNPs with r2 >= 0.1 (in windows of 50 SNPs and steps of 5 SNPs).

To test for admixture, we computed the f3 statistics of the form f3(CONVERGE; ref1, ref2), iterating through all possible pairs of reference populations found on the Human Origin Array data. A significantly negative value of f3 is suggestive of admixture. We further corroborated the observed evidence of admixture by computing the D statistics of the form D(Mbuti, X, CONVERGE, Metropolitan Chinese). Here, X is a potential admixing source population of interest from the Human Origin Array data, while CONVERGE is a specific CONVERGE province tested. The Metropolitan Chinese population is based on a random sample of 200 CONVERGE individuals from the four municipalities in China (Tianjin, Beijing, Shanghai, Chongqing), used as representative of an average Chinese (these four populations are themselves not tested for admixture). Therefore, this D statistic is testing localized signals of admixture by excess of allele sharing between X and CONVERGE province, compared to the metropolitan Chinese individuals. To evaluate shared history between ancient populations and CONVERGE population as measured by allele frequency covariances, we used the outgroup f3 statistics of the form f3(Outgroup; CONVERGE, Ancient), where the Human Origin Array Mbuti population was used as the outgroup. We used qp3Pop and qpDstat package implemented in ADMIXTOOLS (v3.0). No further LD pruning was done as the statistical significance was assessed using the default blocked jackknife approach implemented in ADMIXTOOLS.

For admixture signals from Northern Europeans into Han Chinese from Northwestern China, which more likely reflects a one-pulse admixture model (rather than isolation by distance model among nearby populations), we used ALDER v1.03 (Loh et al. 2013) to estimate the date of the admixture and an f4 ratio test to estimate the admixture proportions (qpF4ratio in ADMIXTOOLS) (Patterson et al. 2012). For ALDER, we looked for successful fit of LD-decay curves corresponding to a pair of Northern European and Southern Chinese population serving as the proxies of ancestral populations to the Northwestern provinces (Gansu, Shaanxi, Shanxi) where we detected admixture signals. We report the fit with largest amplitude from fitting of exponential curves from two source populations that correspond to similar geographical regions as the two source populations reported by f3 test (as the exact two source populations may not produce a successful fit). For f4 ratio test, we used the two source populations inferred from f3 test or from ALDER, and the Sardinians and Chimp sequences as the outgroups to European source population and human populations, respectively. The Sardinians were used as an outgroup of choice because Southern Europeans in general did not contribute as prominently to the admixture signals we detected among Han Chinese, and that the isolated Sardinians more likely reflect the genetic ancestry of ancient Southern Europe or Near East Anatolia (Chiang et al. 2016). Replacing Sardinians with Sicilians or Early Neolithic Farmer samples did not significantly change the results.

### Estimating Neandertal ancestry across CONVERGE

As a simple way to assess the Neanderthal ancestry, for a target population t in the CONVERGE dataset, we computed the following statistic (analogous to the statistic proposed in (Mallick et al. 2016)). We define a set of SNPs, *nd*10, where all 43 African genomes from 17 population in the Simons Genome Diversity Project (SGDP) are ancestral (*i.e.*, match the human ancestor), a randomly picked allele from the Altai Neanderthal (Prufer et al. 2014) is derived while a randomly picked allele from the high-coverage Denisovan individual (Meyer et al. 2012) is ancestral and not all genotypes in t are missing. We then computed an estimate of Neanderthal ancestry in population t:

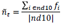

Here *f*_*i*_ is the derived allele frequency at SNP i in population t. We converted this into an estimator of Neanderthal ancestry by rescaling the statistics so that population from Beijing are assigned an ancestry of 1.89% matching previous estimates (Prufer et al. 2014).

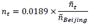

*n*_*t*_ is not an unbiased estimate of Neanderthal ancestry. Nevertheless, differences in these statistics are informative of differential relationships to the Neanderthals. We computed differences in *n*_*t*_ across populations using the individuals from Beijing as baseline. We assessed statistical significance using a block jackknife with 10 Mb blocks. **Table S6** shows that, overall, most populations have a similar relationship to the Neanderthals as the population from Beijing.

### PCA-based Selection Scan

We considered only the top 2 PCs in CONVERGE as these reflect population structure that are interpretable with respect to geography. We initially screened approximately 429,935 randomly chosen SNPs with MAF > 10% in CONVERGE, using a genome-wide significance threshold of 5.8x10-8 (corresponding roughly to Bonferroni correction for testing 430K SNPs and 2 PCs), and then followed up each significant locus by a fine-mapping strategy that uses all SNPs available in CONVERGE with MAF > 1% in order to narrow the list of possible causal variant. PCA-based selection statistics was calculated using the “pcaselection” program in Eigenstrat (v6.1.1) (Galinsky et al. 2016).

As examples of adaptive introgression from archaic humans have been noted previously (Racimo et al. 2017), we examined whether any of the loci detected here may be due to archaic human ancestry. We examined two previously described summary statistics based on uniquely shared sites with archaic humans (Racimo et al. 2017) and compared the statistics at loci detected using PCA-based selection scan against the genomic background. Specifically, for the first statistic we divided the genome into non-overlapping windows of 40kb, and for each window tallied the number of sites where the archaic genome is homozygous for the derived allele and where the derived allele is low frequency (< 1%) in 1000 Genomes YRI but high frequency (> 20%) in CONVERGE (the “uniquely shared sites”). For the second statistic, we computed the 95% quantile of derived allele frequencies in CONVERGE for the uniquely shared sites in each window. A window size of 40kb was used because that is the mean length of introgressed haplotype from archaic humans (Prufer et al. 2014; Racimo et al. 2017). We then ranked each window according to these two statistics and examined the rank of the window containing the lead SNP from PCA-based selection scan. We find no supporting evidence that any of the detected loci shows extreme distribution in these two summary statistics, with exception of the signal on chr20 in PC1, where the window around it ranked in the top 3.6 and 1.0 percentile for the two statistics when using Altai Neandertal, or top 3.8 and 1.0 percentile when using Vindija Neandertal.

## Data Access

## Acknowledgements

C.W.K.C. was supported in part by T32NS048004 from National Institute of Neurological Disorders and Strokes. C.R.R. and S.S. were supported in part by 4R00GM111744 from the National Institute of Health. C.R.R. was also supported by the NIH Training Grant T32HG002536 in Genomic Analysis and Interpretation. W.W.K. was supported by the Wellcome Trust (WT097307). N.C. was supported by an ESPOD Fellowship from EMBL-EBI and Wellcome Trust Sanger Institute.

## References

1000 Genomes Project C, Auton A, Brooks LD, Durbin RM, Garrison EP, Kang HM, Korbel JO, Marchini JL, McCarthy S, McVean GA et al. 2015. A global reference for human genetic variation. Nature 526: 68–74.

Barreiro LB, Laval G, Quach H, Patin E, Quintana-Murci L. 2008. Natural selection has driven population differentiation in modern humans. Nat Genet 40: 340–345.

Brooks PJ, Enoch MA, Goldman D, Li TK, Yokoyama A. 2009. The alcohol flushing response: an unrecognized risk factor for esophageal cancer from alcohol consumption. PLoS Med 6: e50.

Cai N, Bigdeli TB, Kretzschmar WW, Li Y, Liang J, Hu J, Peterson RE, Bacanu S, Webb BT, Riley B et al. 2017. 11,670 whole-genome sequences representative of the Han Chinese population from the CONVERGE project. Sci Data 4: 170011.

Chen CH, Yang JH, Chiang CWK, Hsiung CN, Wu PE, Chang LC, Chu HW, Chang J, Song IW, Yang SL et al. 2016. Population structure of Han Chinese in the modern Taiwanese population based on 10,000 participants in the Taiwan Biobank project. Hum Mol Genet 25: 5321–5331.

Chen J, Zheng H, Bei JX, Sun L, Jia WH, Li T, Zhang F, Seielstad M, Zeng YX, Zhang X et al. 2009. Genetic structure of the Han Chinese population revealed by genome--wide SNP variation. Am J Hum Genet 85: 775–785.

Chiang CW, Marcus JH, Sidore C, Al-Asadi H, Zoledziewska M, Pitzalis M, Busonero F, Maschio A, Pistis G, Steri M et al. 2016. Population history of the Sardinian people inferred from whole-genome sequencing. bioRxiv DOI: https://doi.org/10.1101/092148.

Cockburn IA, Mackinnon MJ, O’Donnell A, Allen SJ, Moulds JM, Baisor M, Bockarie M, Reeder JC, Rowe JA. 2004. A human complement receptor 1 polymorphism that reduces Plasmodium falciparum rosetting confers protection against severe malaria. Proc Natl Acad Sci U S A 101: 272–277.

Converge C. 2015. Sparse whole-genome sequencing identifies two loci for major depressive disorder. Nature 523: 588–591.

Frosst P, Blom HJ, Milos R, Goyette P, Sheppard CA, Matthews RG, Boers GJ, den Heijer M, Kluijtmans LA, van den Heuvel LP et al. 1995. A candidate genetic risk factor for vascular disease: a common mutation in methylenetetrahydrofolate reductase. Nat Genet 10: 111–113.

Fu Q, Posth C, Hajdinjak M, Petr M, Mallick S, Fernandes D, Furtwangler A, Haak W, Meyer M, Mittnik A et al. 2016. The genetic history of Ice Age Europe. Nature 534: 200–205.

Fumagalli M, Moltke I, Grarup N, Racimo F, Bjerregaard P, Jorgensen ME, Korneliussen TS, Gerbault P, Skotte L, Linneberg A et al. 2015. Greenlandic Inuit show genetic signatures of diet and climate adaptation. Science 349: 1343–1347.

Galinsky KJ, Bhatia G, Loh PR, Georgiev S, Mukherjee S, Patterson NJ, Price AL. 2016. Fast Principal-Component Analysis Reveals Convergent Evolution of ADH1B in Europe and East Asia. Am J Hum Genet 98: 45–472.

Genome of the Netherlands C. 2014. Whole-genome sequence variation, population structure and demographic history of the Dutch population. Nat Genet 46: 818–825.

Grossman SR, Andersen KG, Shlyakhter I, Tabrizi S, Winnicki S, Yen A, Park DJ, Griesemer D, Karlsson EK, Wong SH et al. 2013. Identifying recent adaptations in large-scale genomic data. Cell 152: 703–713.

Gudbjartsson DF, Helgason H, Gudjonsson SA, Zink F, Oddson A, Gylfason A, Besenbacher S, Magnusson G, Halldorsson BV, Hjartarson E et al. 2015. Large-scale whole-genome sequencing of the Icelandic population. Nat Genet 47: 435–444.

Haak W, Lazaridis I, Patterson N, Rohland N, Mallick S, Llamas B, Brandt G, Nordenfelt S, Harney E, Stewardson K et al. 2015. Massive migration from the steppe was a source for Indo-European languages in Europe. Nature 522: 207–211.

Han Y, Gu S, Oota H, Osier MV, Pakstis AJ, Speed WC, Kidd JR, Kidd KK. 2007. Evidence of positive selection on a class I ADH locus. Am J Hum Genet 80: 441–456.

Harris K, Nielsen R. 2016. The Genetic Cost of Neanderthal Introgression. Genetics 203: 881–891.

Hedlund GL, Longo N, Pasquali M. 2006. Glutaric acidemia type 1. Am J Med Genet C Semin Med Genet 142C: 86–94.

Hellenthal G, Busby GB, Band G, Wilson JF, Capelli C, Falush D, Myers S. 2014. A genetic atlas of human admixture history. Science 343: 747–751.

Higasa K, Kukita Y, Kato K, Wake N, Tahira T, Hayashi K. 2009. Evaluation of haplotype inference using definitive haplotype data obtained from complete hydatidiform moles, and its significance for the analyses of positively selected regions. PLoS Genet 5: e1000468.

Hirayasu K, Ohashi J, Tanaka H, Kashiwase K, Ogawa A, Takanashi M, Satake M, Jia GJ, Chimge NO, Sideltseva EW et al. 2008. Evidence for natural selection on leukocyte immunoglobulin-like receptors for HLA class I in Northeast Asians. Am J Hum Genet 82: 1075–1083.

Ho P-T. 1976. The Cradle of the East: An Inquiry into the Indigenous Origins of Techniques and Ideas of Neolithic and Early Historic China, 5000-1000 B.C. The University of Chicago Press, Chicago.

Jablonski NG, Chaplin G. 2010. Colloquium paper: human skin pigmentation as an adaptation to UV radiation. Proc Natl Acad Sci U S A 107 Suppl 2: 8962–8968.

Jin ZB, Huang XF, Lv JN, Xiang L, Li DQ, Chen J, Huang C, Wu J, Lu F, Qu J. 2014. SLC7A14 linked to autosomal recessive retinitis pigmentosa. Nat Commun 5: 3517.

Jones MK, Liu X. 2009. Archaeology. Origins of agriculture in East Asia. Science 324: 730–731.

Juric I, Aeschbacher S, Coop G. 2016. The Strength of Selection against Neanderthal Introgression. PLoS Genet 12: e1006340.

Kim BY, Lohmueller KE. 2015. Selection and reduced population size cannot explain higher amounts of Neandertal ancestry in East Asian than in European human populations. Am J Hum Genet 96: 454–461.

Lawson DJ, Hellenthal G, Myers S, Falush D. 2012. Inference of population structure using dense haplotype data. PLoS Genet 8: e1002453.

Lazaridis I, Nadel D, Rollefson G, Merrett DC, Rohland N, Mallick S, Fernandes D, Novak M, Gamarra B, Sirak K et al. 2016. Genomic insights into the origin of farming in the ancient Near East. Nature 536: 419–424.

Lazaridis I Patterson N Mittnik A Renaud G Mallick S Kirsanow K Sudmant PH Schraiber JG Castellano S Lipson M et al. 2014. Ancient human genomes suggest three ancestral populations for present-day Europeans. Nature 513: 409–413.

Lek M, Karczewski KJ, Minikel EV, Samocha KE, Banks E, Fennell T, O’Donnell-Luria AH, Ware JS, Hill AJ, Cummings BB et al. 2016. Analysis of protein-coding genetic variation in 60,706 humans. Nature 536: 285–291.

Li H, Gu S, Han Y, Xu Z, Pakstis AJ, Jin L, Kidd JR, Kidd KK. 2011a. Diversification of the ADH1B gene during expansion of modern humans. Ann Hum Genet 75: 497–507.

Li Y, Sidore C, Kang HM, Boehnke M, Abecasis GR. 2011b. Low-coverage sequencing: implications for design of complex trait association studies. Genome Res 21: 940–951.

Liu X, Ong RT, Pillai EN, Elzein AM, Small KS, Clark TG, Kwiatkowski DP, Teo YY. 2013. Detecting and characterizing genomic signatures of positive selection in global populations. Am J Hum Genet 92: 866–881.

Loh PR, Lipson M, Patterson N, Moorjani P, Pickrell JK, Reich D, Berger B. 2013. Inferring admixture histories of human populations using linkage disequilibrium. Genetics 193: 1233–1254.

Mallick S, Li H, Lipson M, Mathieson I, Gymrek M, Racimo F, Zhao M, Chennagiri N, Nordenfelt S, Tandon A et al. 2016. The Simons Genome Diversity Project: 300 genomes from 142 diverse populations. Nature 538: 201–206.

Manichaikul A, Mychaleckyj JC, Rich SS, Daly K, Sale M, Chen WM. 2010. Robust relationship inference in genome-wide association studies. Bioinformatics 26: 2867–2873.

Mathieson I, Lazaridis I, Rohland N, Mallick S, Patterson N, Roodenberg SA, Harney E, Stewardson K, Fernandes D, Novak M et al. 2015. Genome-wide patterns of selection in 230 ancient Eurasians. Nature 528: 499–503.

McKay JD Truong T Gaborieau V Chabrier A Chuang SC Byrnes G Zaridze D Shangina O Szeszenia-Dabrowska N Lissowska J et al. 2011. A genome-wide association study of upper aerodigestive tract cancers conducted within the INHANCE consortium. PLoS Genet 7: e1001333.

McLaren W, Gil L, Hunt SE, Riat HS, Ritchie GR, Thormann A, Flicek P, Cunningham F. 2016. The Ensembl Variant Effect Predictor. Genome Biol 17: 122.

Meyer M, Kircher M, Gansauge MT, Li H, Racimo F, Mallick S, Schraiber JG, Jay F, Prufer K, de Filippo C et al. 2012. A high-coverage genome sequence from an archaic Denisovan individual. Science 338: 222–226.

Moorjani P, Sankararaman S, Fu Q, Przeworski M, Patterson N, Reich D. 2016. A genetic method for dating ancient genomes provides a direct estimate of human generation interval in the last 45,000 years. Proc Natl Acad Sci U S A 113: 5652–5657.

Munoz-Moran E, Dieguez-Lucena JL, Fernandez-Arcas N, Peran-Mesa S, Reyes-Engel A. 1998. Genetic selection and folate intake during pregnancy. Lancet 352: 1120–1121.

Naeem Z, Newton V. 1996. Prevalence of sensorineural hearing loss in Asian children. Br J Audiol 30: 332–339.

Oota H, Pakstis AJ, Bonne-Tamir B, Goldman D, Grigorenko E, Kajuna SL, Karoma NJ, Kungulilo S, Lu RB, Odunsi K et al. 2004. The evolution and population genetics of the ALDH2 locus: random genetic drift, selection, and low levels of recombination. Ann Hum Genet 68: 93–109.

Pare G, Chasman DI, Parker AN, Zee RR, Malarstig A, Seedorf U, Collins R, Watkins H, Hamsten A, Miletich JP et al. 2009. Novel associations of CPS1, MUT, NOX4, and DPEP1 with plasma homocysteine in a healthy population: a genome-wide evaluation of 13 974 participants in the Women’s Genome Health Study. Circ Cardiovasc Genet 2: 142–150.

Park BL, Kim JW, Cheong HS, Kim LH, Lee BC, Seo CH, Kang TC, Nam YW, Kim GB, Shin HD et al. 2013. Extended genetic effects of ADH cluster genes on the risk of alcohol dependence: from GWAS to replication. Hum Genet 132: 657–668.

Patterson N, Moorjani P, Luo Y, Mallick S, Rohland N, Zhan Y, Genschoreck T, Webster T, Reich D. 2012. Ancient admixture in human history. Genetics 192: 1065–1093.

Peng Y, Shi H, Qi XB, Xiao CJ, Zhong H, Ma RL, Su B. 2010. The ADH1B Arg47His polymorphism in east Asian populations and expansion of rice domestication in history. BMC Evol Biol 10: 15.

Peter BM. 2016. Admixture, Population Structure, and F-Statistics. Genetics 202: 1485–1501.

Price AL, Weale ME, Patterson N, Myers SR, Need AC, Shianna KV, Ge D, Rotter JI, Torres E, Taylor KD et al. 2008. Long-range LD can confound genome scans in admixed populations. Am J Hum Genet 83: 132–135; author reply 135-139.

Prufer K, Racimo F, Patterson N, Jay F, Sankararaman S, Sawyer S, Heinze A, Renaud G, Sudmant PH, de Filippo C et al. 2014. The complete genome sequence of a Neanderthal from the Altai Mountains. Nature 505: 43–49.

Racimo F, Gokhman D, Fumagalli M, Ko A, Hansen T, Moltke I, Albrechtsen A, Carmel L, Huerta-Sanchez E, Nielsen R. 2017. Archaic Adaptive Introgression in TBX15/WARS2. Mol Biol Evol 34: 509–524.

Raghavan M, Skoglund P, Graf KE, Metspalu M, Albrechtsen A, Moltke I, Rasmussen S, Stafford TW, Jr., Orlando L, Metspalu E et al. 2014. Upper Palaeolithic Siberian genome reveals dual ancestry of Native Americans. Nature 505: 87–91.

Rosenberg N, Murata M, Ikeda Y, Opare-Sem O, Zivelin A, Geffen E, Seligsohn U. 2002. The frequent 5,10-methylenetetrahydrofolate reductase C677T polymorphism is associated with a common haplotype in whites, Japanese, and Africans. Am J Hum Genet 70: 758–762.

Sankararaman S, Mallick S, Dannemann M, Prufer K, Kelso J, Paabo S, Patterson N, Reich D. 2014. The genomic landscape of Neanderthal ancestry in present-day humans. Nature 507: 354–357.

Sankararaman S, Mallick S, Patterson N, Reich D. 2016. The Combined Landscape of Denisovan and Neanderthal Ancestry in Present-Day Humans. Curr Biol 26: 1241–1247.

Sidore C, Busonero F, Maschio A, Porcu E, Naitza S, Zoledziewska M, Mulas A, Pistis G, Steri M, Danjou F et al. 2015. Genome sequencing elucidates Sardinian genetic architecture and augments association analyses for lipid and blood inflammatory markers. Nat Genet 47: 1272–1281.

Simonti CN, Vernot B, Bastarache L, Bottinger E, Carrell DS, Chisholm RL, Crosslin DR, Hebbring SJ, Jarvik GP, Kullo IJ et al. 2016. The phenotypic legacy of admixture between modern humans and Neandertals. Science 351: 737–741.

Suo C, Xu H, Khor CC, Ong RT, Sim X, Chen J, Tay WT, Sim KS, Zeng YX, Zhang X et al. 2012. Natural positive selection and north-south genetic diversity in East Asia. Eur J Hum Genet 20: 102–110.

UK10K C, Walter K, Min JL, Huang J, Crooks L, Memari Y, McCarthy S, Perry JR, Xu C, Futema M et al. 2015. The UK10K project identifies rare variants in health and disease. Nature 526: 82–90.

van Meurs JB, Pare G, Schwartz SM, Hazra A, Tanaka T, Vermeulen SH, Cotlarciuc I, Yuan X, Malarstig A, Bandinelli S et al. 2013. Common genetic loci influencing plasma homocysteine concentrations and their effect on risk of coronary artery disease. Am J Clin Nutr 98: 668–676.

Vernot B, Akey JM. 2014. Resurrecting surviving Neandertal lineages from modern human genomes. Science 343: 1017–1021.

Vernot B, Akey JM. 2015. Complex history of admixture between modern humans and Neandertals. Am J Hum Genet 96: 448–453.

Welter D, MacArthur J, Morales J, Burdett T, Hall P, Junkins H, Klemm A, Flicek P, Manolio T, Hindorff L et al. 2014. The NHGRI GWAS Catalog, a curated resource of SNP--trait associations. Nucleic Acids Res 42: D1001–1006.

Wen B, Li H, Lu D, Song X, Zhang F, He Y, Li F, Gao Y, Mao X, Zhang L et al. 2004. Genetic evidence supports demic diffusion of Han culture. Nature 431: 302–305.

Whiffin N, Minikel E, Walsh R, O’Donnell-Luria AH, Karczewski K, Ing AY, Barton PJR, Funke B, Cook SA, MacArthur D et al. 2017. Using high-resolution variant frequencies to empower clinical genome interpretation. Genet Med DOI:10.1038/gim.2017.26.

Xu S, Yin X, Li S, Jin W, Lou H, Yang L, Gong X, Wang H, Shen Y, Pan X et al. 2009. Genomic dissection of population substructure of Han Chinese and its implication in association studies. Am J Hum Genet 85: 762–774.

Yafei W, Lijun P, Jinfeng W, Xiaoying Z. 2012. Is the prevalence of MTHFR C677T polymorphism associated with ultraviolet radiation in Eurasia? J Hum Genet 57: 780–786.

Yang J, Jin ZB, Chen J, Huang XF, Li XM, Liang YB, Mao JY, Chen X, Zheng Z, Bakshi A et al. 2017. Genetic signatures of high-altitude adaptation in Tibetans. Proc Natl Acad Sci U S A 114: 4189–4194.

Yang J, Lee SH, Goddard ME, Visscher PM. 2011. GCTA: a tool for genome-wide complex trait analysis. Am J Hum Genet 88: 76–82.

Ye K, Gao F, Wang D, Bar-Yosef O, Keinan A. 2017. Dietary adaptation of FADS genes in Europe varied across time and geography. Natue Ecology and Evolution 1: 0167.

You QS, Xu L, Wang YX, Liang QF, Cui TT, Yang XH, Wang S, Yang H, Jonas JB. 2013. Prevalence of retinitis pigmentosa in North China: the Beijing Eye Public Health Care Project. Acta Ophthalmol 91: e499–500.

Zhao YB, Zhang Y, Zhang QC, Li HJ, Cui YQ, Xu Z, Jin L, Zhou H, Zhu H. 2015. Ancient DNA reveals that the genetic structure of the northern Han Chinese was shaped prior to 3,000 years ago. PLoS One 10: e0125676.

Zhao Z. 2011. New Archaeobotanic Data for the Study of the Origins of Agriculture in China. Current Anthropology 52: S295–S306.

